# Robust and accurate method for measuring tumor volume using optical 3D scanning for nonclinical in vivo efficacy study

**DOI:** 10.1101/2024.05.07.592924

**Authors:** Takuma Kobayashi, Mayumi Katsumata, Yoshiki Nakamura, Yuri Terado, Hideo Araki, Eiki Maeda

## Abstract

In a nonclinical *in vivo* efficacy test for anticancer drugs, immunodeficient mice subcutaneously transplanted with human cancer cells were quantified and evaluated with regard to the manner in which the skin bulges where locally proliferated cancer cells regress after drug administration. A caliper is conventionally used to measure the tumor bulge. However, its volume is an estimated value and results in high variability. Alternatively, cancer cell lines that express genetically encoded marker genes have been used in recent years for optical and nondestructive measurements. However, estimations using calipers exhibit large errors, and biological tissues have low light transparency. This hinders quantitative optical measurements. In addition, variations in measurements owing to subjective and human operations are likely.

From the chemistry, manufacturing, and control (CMC) perspective, precise measurement is required to evaluate drug efficacy and quality. Therefore, we aimed to eliminate errors caused by the use of estimated values, subjectivity, and human manipulation by precisely quantifying the volume of the tumor bulge using a 3D scanner.

This study demonstrated that optical 3D scanner measurements were accurate, had low variability, and was highly correlated with tumor weight. The tumor bulge was observed to vary to a flattened oval dome shape rather than a semicircle. This caused high variability in measurements of tumor volume. However, the proposed 3D scanner was more sensitive to volumetric regression than the caliper. Additionally, it exhibited drug efficacies with higher resolution than the caliper. Furthermore, the high linearity of the scanner provided more accurate measurements over a wider range of tumor sizes than luminescence imaging. The accurate and sensitive properties of such 3D scanners are also likely to make these exceptionally effective analytical tools for ensuring product equivalency when modifying raw materials or manufacturing processes in the development of cell therapy products.

As described above, robust and accurate drug efficacy measurements using nondestructive and noninvasive 3D scanners that require no training and are convenient to operate provide many analytical improvements and advantages. This is likely to play an important role in 1) the efficacy evaluation of cell therapy products that have large variations originating from the raw materials and large differences between manufacturing lots and 2) the quality evaluation, property analysis of the characteristics of variations in the shape of tumor bulges over time, and comparability testing of the products in the CMC section of pharmaceutical companies.

## Introduction

To test the *in vivo* efficacy of anticancer drugs, xenograft models (in which human cancer cells are transplanted into immunocompromised mice) are generally used in nonclinical trials (Teicher, 2006; Sausville and Burger, 2006; Ruggeri, et al., 2013; Liu et al., 2023). In a xenograft model in which human tumor cells are transplanted subcutaneously (Fig. 1a), the transplanted cancer cells proliferate locally and bulge (Fig. 1b). Therefore, after administering anticancer drugs to these tumor-bearing mice, the growth or regression of tumor cells at the bulge site can be measured externally and quantified reproducibly to evaluate their efficacy.

**Figure 1.**
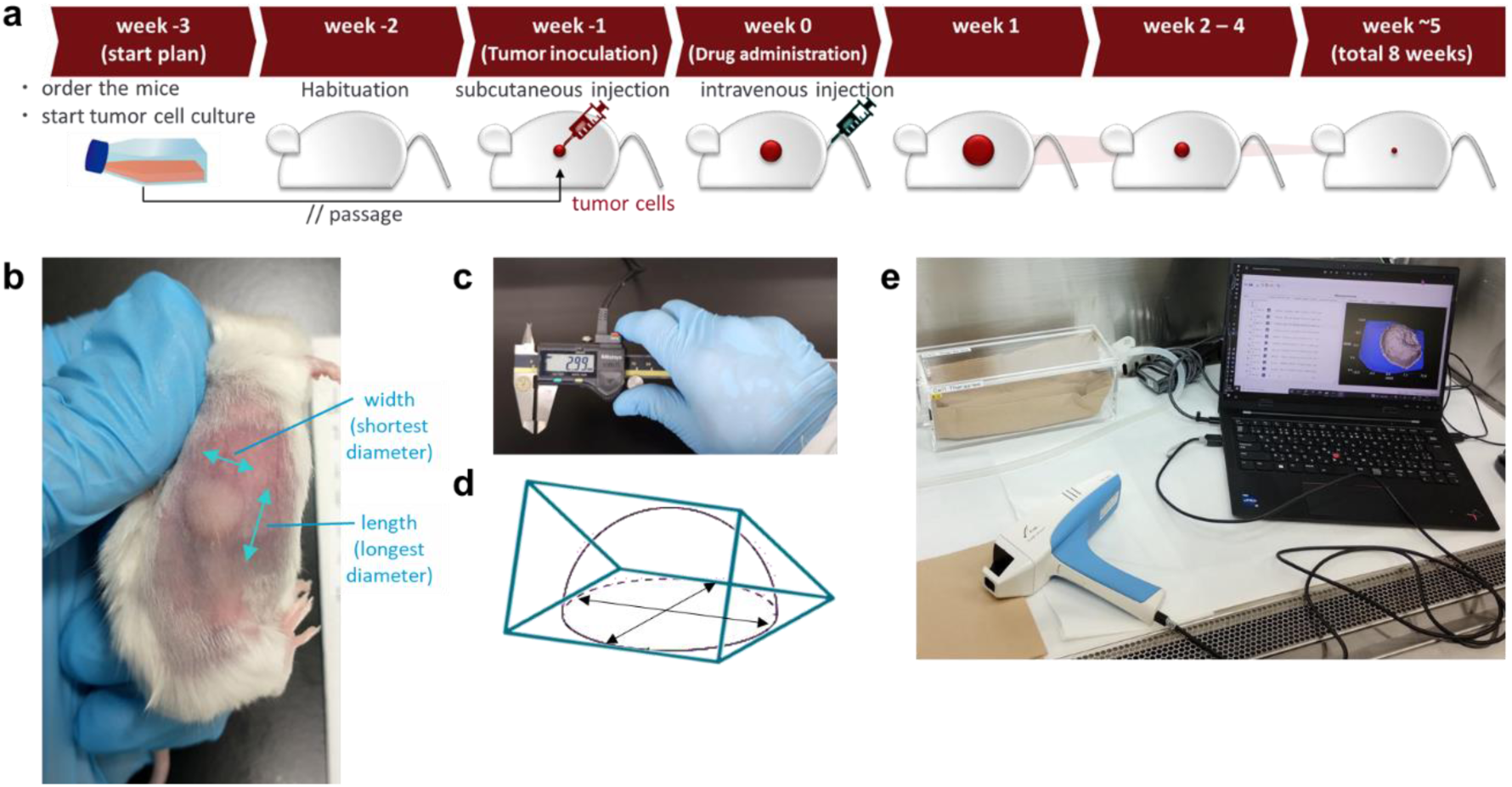
A schematic image of subcutaneous xenograft in vivo study and measurement methods for tumor volume. (a) Schematic showing a typical subcutaneous xenograft model. Cultured human tumor cells were injected subcutaneously into immunodeficient mice. After inoculation, the transplanted tumor cells gradually proliferated and formed protrusions on the skin of the mice, as shown in (b). Next, the mice were intravenously administered cell therapy drugs via the tail vein. After drug administration, the transplanted tumor cells were expected to regress gradually. This process was monitored over time by measuring their size using calipers and luminescence. (b) Example of the bulge of a subcutaneous tumor in a mouse. The bulge was measured by subjectively determining the longest and shortest diameters (major and minor axes) of the distorted elliptical dome structure. (c) Tumor size measured using digital calipers. (d) Image of the actual tumor protrusion and estimated triangular prism. (e) Typical 3D scanner set.

Tumor volume is conventionally estimated by measuring the shortest (width) and longest (length) diameters of the elliptical dome-shaped tumor bulge non-invasively and over time (Fig. 1b) using calipers (Fig. 1c) and applying the following formula: Volume = (width)^2 × length / 2 [mm^3] (Ovejera, et al., 1978). However, the value calculated using this formula is that of a triangular pyramid, which is different from the actual tumor volume with an elliptical hemispherical dome shape (Fig. 1d). Tumor bulges are occasionally irregularly shaped. This hinders the accurate measurement of their volumes using calipers. Furthermore, the criteria for determining the location, width, and length of the tumor bulge may vary among operators. In addition, the shape of the tumor bulge may vary depending on the holding posture and the amount of force exerted on the skin of the mouse. The inclusion of many subjective and artificial manipulations can result in different measurement values and variability. These subjective and artificial manipulations result in large variations in the measurements. For the above reasons, caliper operation requires prior preparation such as practice to become proficient in handling the caliper and mice and to match measurements with subjective values so that measurements do not differ among operators (Table 1).

**Table 1.** List of steps and time required for each measurement method. Anesthetization of the mouse was optional. However, it was recommended to improve the accuracy of the measurement values.

| Measurement using caliper | time | Luminescence imaging | time | 3D scanning | time |
| --- | --- | --- | --- | --- | --- |
| ↓ [ Preparation ]<br>Practice for matching subjective assessment criteria that differ depending on the operator and measuring with the caliper | ( 2 months ) | [ Preparation ]<br>- Obtaining tumor cells expressing Luciferase<br>- Preliminary experiments such as considering luminescence measurement range and verifying kinetics | ( 2 – 6 months ) | ( No advance preparation required ) | ( 0 ) |
| ↓ [ Optional ] anesthetize the mouse | ( 3 min. ) | anesthetize the mouse | 3 min. | [ Optional ] anesthetize the mouse | ( 3 min. ) |
| ↓ Hold the mouse or lay the mouse down with the tumor-bearing area facing up | 2 s. | Fill the syringe with the prepared Luciferin reagent and Inject Luciferin intraperitoneally | 5 s. | Hold the mouse or lay the mouse down with the tumor-bearing area facing up | 2 s. |
| ↓ Wipe the skin above the tumor-bearing area with ethanol cotton (to improve visibility) | 2 s. | Wait until Luciferin circulates in the mouse body | 10 min / 5 mice | scan with scanner | 1 s. |
| ↓ Determine the longest and shortest diameter of the tumor bulge | 2 s. | Lay the mouse in the imaging box with the tumor-bearing area facing up, and set the mouse on the anesthesia mask | 5 s. |  |  |
| ↓ Measure the longest and shortest diameter using the caliper | 4 s. | Perform the luminescence imaging | 1 s. – 1 min. / 5 mice |  |  |
| ↓ |  | [ Optional ] checking the success or failure of intraperitoneal administration of Luciferin by performing fluorescence imaging using the IVISbrite and analyzing the fluorescence intensity* (* see also Discussion for detail) | ( 15 min. / 5 mice ) |  |  |
| **Total time for typical routine measurements per mouse** | **10 s.** |  | **5 min. 10 s. – 5 min. 22 s.** |  | **3 s.** |

Recently, tumor volume has been measured optically using tumor cell lines expressing genetically encoded fluorescent proteins or oxidative enzymes that produce bioluminescence (Hoffman, 2005; Jong, et al., 2014; Walsh and Quail, 2023). For example, a cell line expressing luciferase was transplanted into mice, and its substrate, luciferin, was administered to the mice to emit bioluminescence for imaging (Sato et al., 2004; Kelkar and De, 2012). Biological tissues have low optical transparency to light in the visible spectrum (400–700 nm). This hinders quantitative optical measurements. For efficient optical imaging, it is preferable to use a “biological optical window” (approximately 700–1400 nm) to circumvent the high absorption range of hemoglobin and water (Kato et al., 1993; Smith, et al., 2009; Hong, et al., 2017; Kobayashi et al., 2016). Long-wavelength, broad-spectrum luminescence has a higher optical transparency through biological tissues than fluorescence, which requires relatively short-wavelength excitation and is generally narrow-spectrum for discrimination from excitation light. In addition, luminescence has a higher signal-to-noise ratio than fluorescence including autofluorescence from living tissues. Accordingly, luminescence with a red-shifted wavelength and broad spectrum tends to be more sensitive to detection than fluorescence at specific wavelengths (Rice et al., 2001; Yeh et al., 2017; Endo and Ozawa, 2020). However, as the tumor tissue enlarges and thickens, the optical transparency decreases, and the tumor tissue obstructs the luminescence from the piled-up tumor cells. Furthermore, to perform luminescence measurements, it is necessary to additionally administer chemicals such as modified synthetic luciferin to tumor cells modified genetically to express luciferase. This operation is expensive and requires training as in the case of caliper operation. The operator should be proficient in safely and reliably injecting the required amount of substrate into the abdominal cavity of the mouse for each measurement, verify the kinetics of luciferin in the experimental system in advance, measure the luminescence at the peak of each time according to the kinetics curve (Fig. 3), maintain the mouse in a fixed position inside the imaging device each time, and measure it using luminescence imaging (Table 1).

As described above, conventional methods such as calipers and luminescence measurements have multiple problems that prevent stringent and precise quantification. In addition, both the measurements require time for training and measurement processes and tend to vary their measurement values because these involve subjective and artificial manipulation. Such variability should be eliminated from the perspective of product quality evaluation and characterization in the chemistry, manufacturing, and control (CMC) of pharmaceutical companies. This is particularly so for the efficacy evaluation of cell therapy products, which involve large variations in raw material origin and whose properties can vary straightforwardly depending on manufacturing lots and storage conditions (Cauchon, et al., 2019). Therefore, we considered that the tumor volume could be quantified stably and precisely without variability caused by estimation, subjectivity, or human manipulation using a 3D scanner with maximum accuracy. The convenience of handling the 3D scanner is also likely to reduce the routine measurement time (Table 1).

In this study, a TM900 (Peira, BELGIUM) was selected as a handheld scanner. It is convenient to operate, is an optical type with a higher spatial resolution than ultrasonic echo or sonar, and does not include estimated values and rather calculates volumes directly from the measured shapes (Fig. 1e). Although several types of optical scanners exist and are used to measure tumor volume (Girit et al., 2008; Mu, et al., 2017; Delgado-SanMartin et al., 2019), few reports precisely compare operator-to-operator differences between scanners and conventional methods such as calipers, luminescence, and tumor weight over time for the same object. Therefore, in this study, we compared the results of these conventional measurement methods and scanners over time and reported the newly revealed characteristics of each method. We report for the first time that this scanner is highly accurate during tumor regression.

## Results

First, the difference in the capability of the caliper and scanner to physically measure tumor bulges of various sizes was investigated. We measured the tumor bulges approximately two–three weeks after subcutaneous injection of GSU cells, a human cell line derived from gastric cancer, into immunodeficient NOD.Cg-*Prkdc*^scid^Il2rg^tm1Wjl^/SzJ (NSG) mice (Fig. 2). By measuring and comparing the same object with caliper and scanner, the simple correlation coefficient (r) was determined to be 0.96. This reveals a high correlation between caliper and scanner measurements for tumors of all sizes (Fig. 2a). Therefore, a scanner can be used rather than a caliper. Subsequently, different operators measured tumor bulges of various sizes using calipers and scanners. The simple correlation coefficient between the values measured by operators A and B using the caliper was r = 0.95 (Fig. 2b), whereas that for the scanner was r = 0.99 (Fig. 2c). In both the results, no bias appeared to exist in the variation depending on specific tumor size. These results illustrate that when using the scanner rather than the caliper, the differences in measurement values between operators are relatively low. Next, to compare the measurement differences between different test populations, the tumor size in the test groups (Gr.1-4 in Fig. 2d, Gr.1-6 in Fig. 2e) of mice administered with different cell therapy products was measured using a caliper or scanner by different operators (Op.A-D in Fig. 2d, Op.A, B in Fig. 2e). Several groups of mice had relatively small tumor sizes, e.g., Gr4 in Fig. 2d and Gr4-6 in Fig. 2e. Because none of the operators had practiced in advance to match the measured values, statistically significant differences existed in the mean values of tumor volumes even within the same group of mice when measured by different operators using a caliper (asterisks in Fig. 2d: ** p = 0.035, *** p = 0.0068). However, no statistically significant differences existed in the mean tumor volumes within the same group of mice when measured using the scanner even when it was by different operators. These results reveal that when using the scanner rather than the caliper, the differences between operators in measurement values are exceptionally small. It is indicated that prior practice is essential to reduce the differences between operators in measurement values when using the caliper correctly. Therefore, measurement with a 3D scanner improves accuracy by eliminating differences between operators. Unlike calipers, advanced practice and preparation is unnecessary. This contributes to a reduction in man-hours. In particular, the tumor volume obtained by caliper measurement was an estimated value, whereas that measured using a 3D scanner was closer to the actual value.

**Figure 2.**
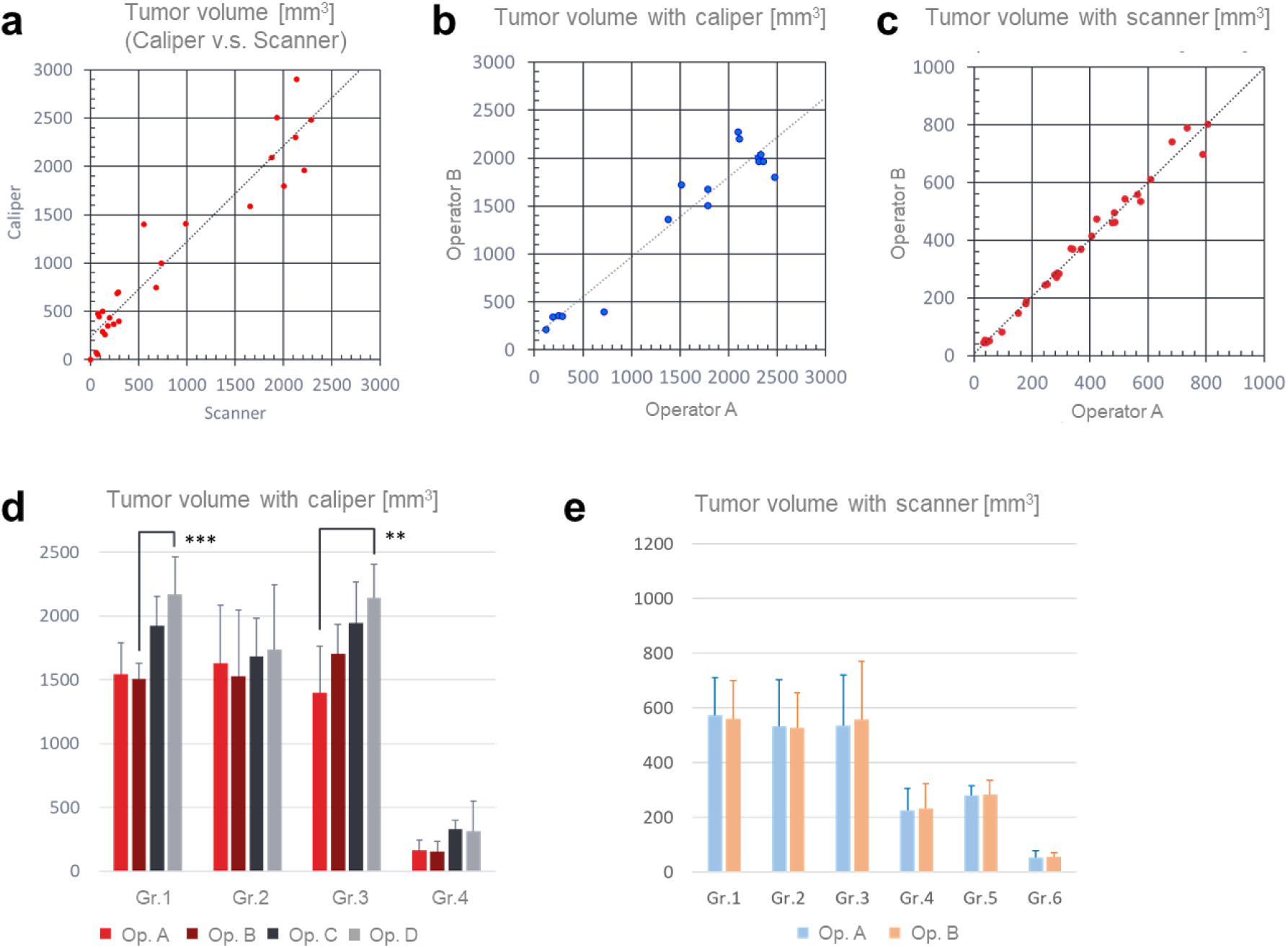
Scanner measurements are more accurate and reproducible than caliper measurements. This contributes to eliminating operator-to-operator errors. After tumor bearing, the tumor volumes were measured in the test group of mice administered with various cell therapy drugs. (a) The graph shows the comparative results of tumor volume measurements of various sizes using a caliper and 3D scanner. The correlation coefficient (r) is 0.96. (b) The graph shows the comparative results of tumor volume measurements of various sizes using calipers by different operators. The operators had no prior training to match their measurements. The average measurement error between operators was 80.6% (n = 16). r = 0.95. (c) Graph showing the comparative results of tumor volume measurements of various sizes by different operators using a 3D scanner. The operators had no prior training to match their measurements. The average measurement error between operators was 6.5% (n = 60). r = 0.99. After tumor bearing, the tumor size in the test groups of mice administered different cell therapy products was measured using a caliper or scanner by different operators. (d) The graph shows a comparison of tumor volumes measured using calipers by different operators in groups (cages) of mice treated with different cell therapy drugs. The operators had no prior training to match their measurements. The data were analyzed using one-way analysis of variance (ANOVA) followed by Bonferroni’s multiple comparisons. The asterisks indicate statistically significant differences in tumor volume measurements of mice from the same test group by different operators (** p < 0.05, ***p < 0.01, and p-values of 0.035 and 0.0068, respectively). The maximum measurement error between operators was 316.5% (Op. = operator, Gr. = test group of mice). (e) The graph shows the comparison results of tumor volumes measured using the 3D scanner by different operators in groups (cages) of mice treated with different cell therapy drugs. The operators had no prior training to match their measurements. There were no statistically significant differences. The maximum measurement error between operators was 42.3% (Op. = operator, Gr. = test group of mice).

We further verified the feasibility of using 3D scanners with an *in vivo* imaging system (IVIS). The *in vivo* drug efficacy was tested using a xenograft model. Prior to the testing, the kinetics of luciferin in our experimental system were verified (Fig. 3). It is well known that enzyme reactions, metabolic times, and sensitivities differ between different tumor cell lines and mouse models. Therefore, prior verification of the experimental system used was required. The results revealed that the luminescence of GSU cells subcutaneously inoculated into NSG mice peaked 10 min after luciferin administration and then, decayed rapidly. However, a small signal was detectable after 4 h (Fig. 3a, b). In our *in vivo* xenograft model, chimeric antigen receptor (CAR) T cells (a cell therapy product) were administered to subcutaneous tumor-bearing mice. Their efficacy was evaluated by measuring the tumor volume and luminescence intensity using a caliper, a scanner, and an IVIS, respectively (Fig. 4a). Compared with the control group with untransduced (UTD) T-cells without CAR gene transduction, the luminescence intensity in the 1E6 CAR-T cell administration group decreased and disappeared after four weeks of IVIS. The variations over time for each measurement using the calipers, scanners, and IVIS are shown in Fig. 4b. According to the scanner measurements, the tumor volume in the 1E6 CAR-T cell administration group attained a value close to zero at the endpoint (middle graph of Fig. 4b). This value was significantly different from the endpoint value in the UTD cell administration group (p = 0.0274, middle graph in Fig. 4b). In contrast, caliper measurements showed that the tumor volume in the 1E6 CAR-T cell administration group was still 1000 mm^3 at the endpoint (left graph of Fig. 4b). Although this value tended to be statistically different from the endpoint value of the UTD administration group (p = 0.0557, left graph of Fig. 4b), the measured value did not vary significantly from day 7. The IVIS values appeared to have attained a plateau after day 12, and the values did not increase. Although the luminescence intensity in the 1E6 CAR-T cell administration group attained zero at the endpoint by IVIS measurement (right graph of Fig. 4b), this value was not significantly different from that of the UTD administration group (p = 0.296, right graph of Fig. 4b). According to the results obtained using the caliper and scanner, the tumor volume in the PBS- or UTD-administered groups on day 12 was approximately 800–1000 mm^3. It then increased further to approximately 1500 mm^3. This difference in the time course between IVIS and calipers or scanners indicates that the enlarged tissue of the tumor bulge may obstruct the transmission of luminescence. Simultaneously, it may be difficult for luciferin to penetrate the enlarged tumor. Notwithstanding the large number of blood vessels, tumors are generally hypoxic and malnourished because of vascular dysfunction (Carmeliet and Jain, 2011). Luciferins in the bloodstream, such as oxygen and nutrients, may have difficulty penetrating large tumor tissues. It is likely that luciferin has low permeability to large tumor tissues and that the accumulation of necrotic cells reduces light permeability. These results indicate that the in vivo efficacy of CAR-T cells is more sensitive when a 3D scanner is used than when a caliper or IVIS is used.

**Figure 3.**
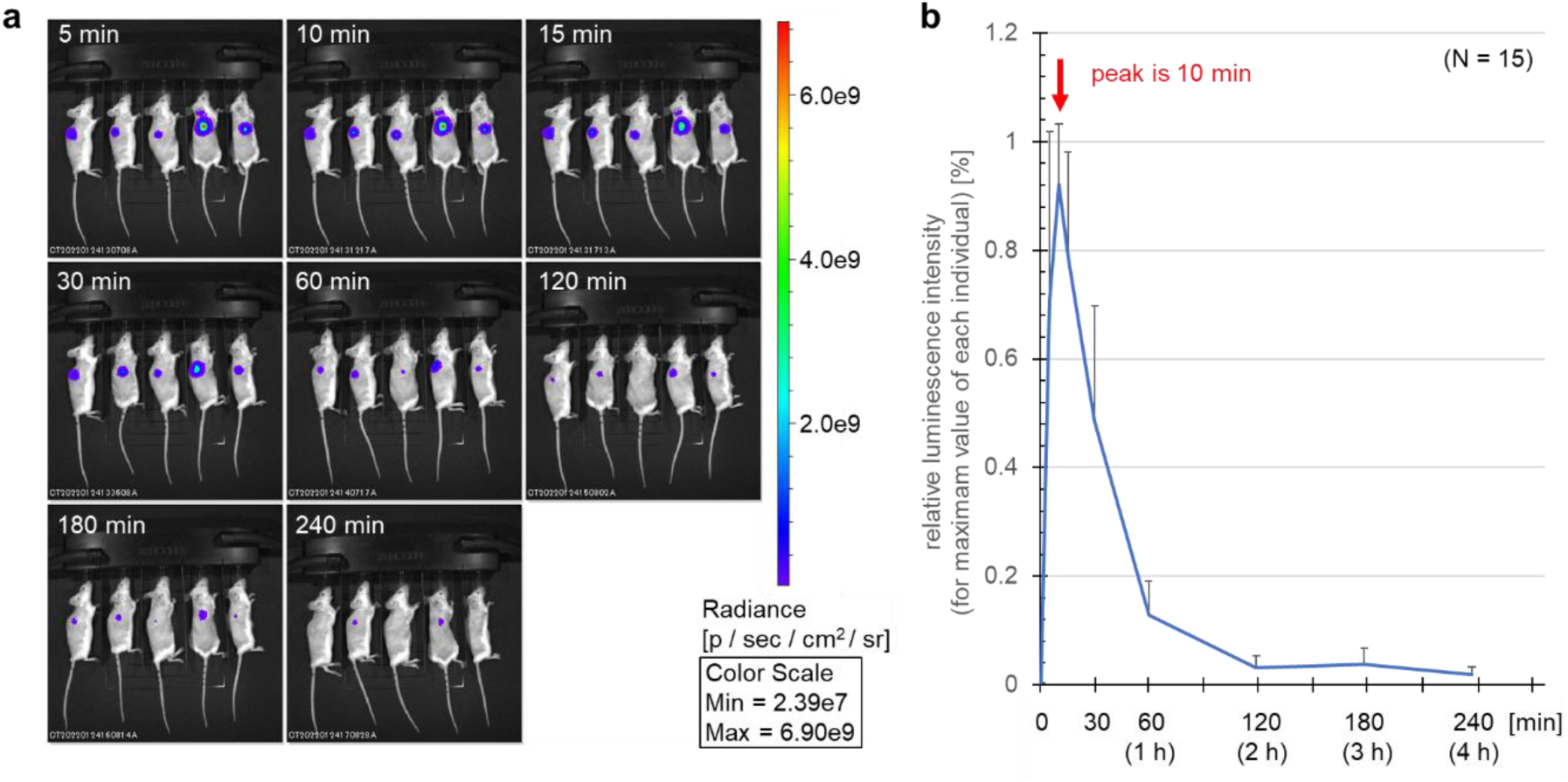
The results of kinetics analysis of D-Luciferin in mice subcutaneously transplanted with 5E6 GSU cells after 1 week. (A) Time-lapse image of luminescence following luciferin administration. The signal intensity is represented in the radiance unit of photons (p) /s/cm^2/steradian (sr). (b) The graph shows the kinetic curve of luciferase activity *in vivo* in 15 mice. The relative luminescence intensity peaked at 10 min.

**Figure 4.**
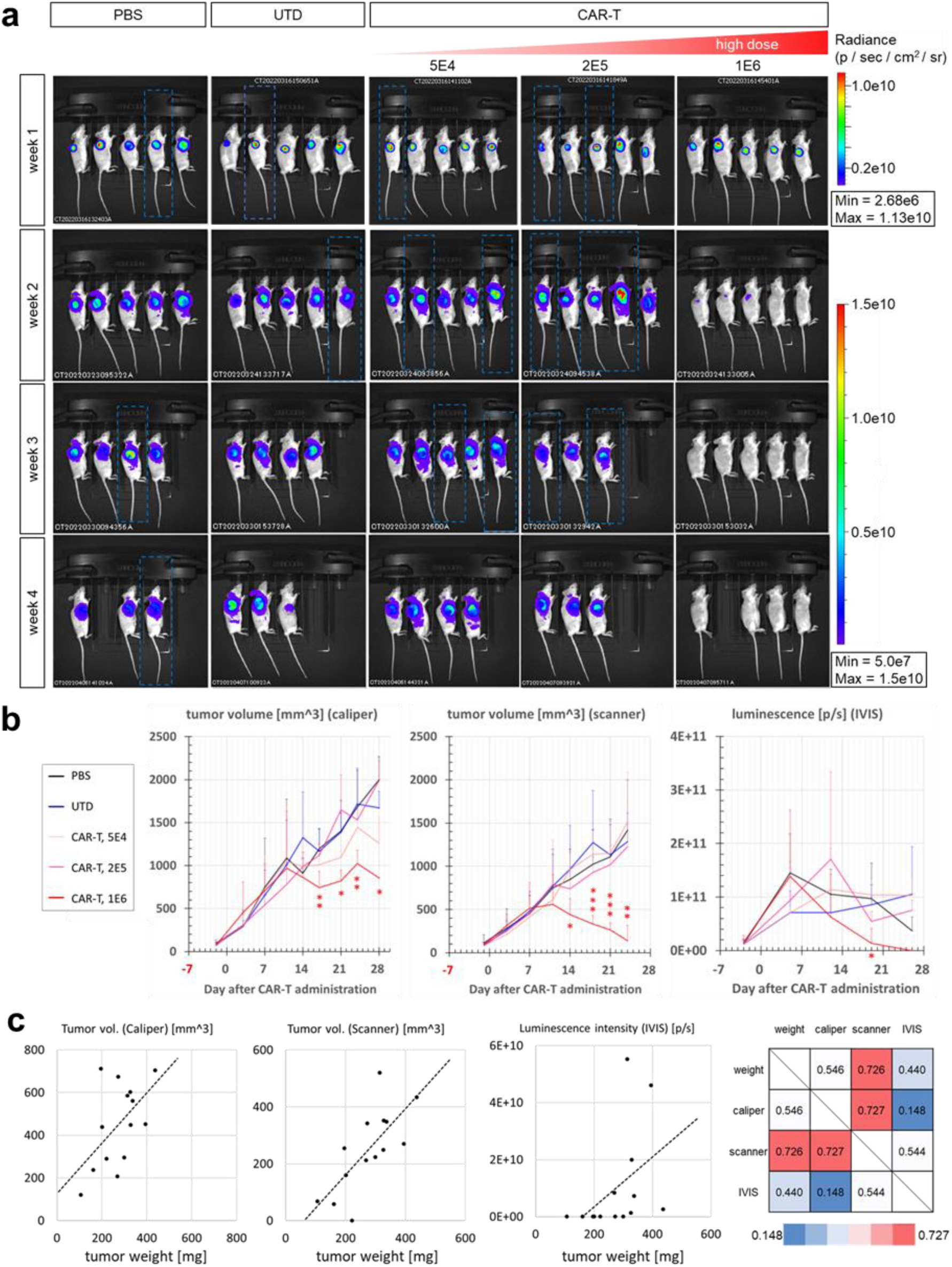
Scanner measurements are more accurate than caliper or IVIS measurements and are highly correlated with tumor weight. This helps eliminate variability. After 5E6 GSU bearing, the tumor volumes were measured using calipers and 3D scanners. Luminescence imaging was performed simultaneously with IVIS over time in the test group of mice administered with various concentrations of cell therapy drugs. (a) Example results of luminescence imaging over time in test groups of mice administered with different numbers of CAR-T cells, UTD, and PBS are shown. The pseudo-color represents the signal intensity of luminescence from the tumor-bearing cells that express luciferase. Mice boxed by the blue broken line are recaptured images obtained by the re-intraperitoneal administration of luciferin. (b) The graphs show the variations in tumor volumes measured using calipers and 3D scanners and the variations in luminescence intensity (photons (p)/s). The data were statistically analyzed using ANOVA followed by Dunnett’s multiple comparisons. The asterisks indicate statistically significant differences between the results of the UTD and various doses of CAR-T cells (*p < 0.1, **p < 0.05, and ***p < 0.01). (c) The graphs show a comparison between tumor weight and volume (measured using a caliper or 3D scanner) and that between tumor weight and luminescence intensity measured using IVIS. The table shows the correlation coefficients for each measurement. The gradual pseudo-colors from blue to red represent the order of increasing correlation coefficients.

Next, we investigated whether the obtained tumor measurement values correlated with the tumor weight (Fig. 4c). The scanner measurements showed the strongest correlation with tumor weight (r = 0.726), whereas the caliper measurements showed a weak correlation with it (r = 0.546). The correlation coefficient between the scanner and caliper measurements was r = 0.727. The IVIS measurements had the largest variance and lowest correlation with tumor weight (r = 0.148). In addition, the luminescence intensity was almost zero when the tumor weight was less than 200–300 mg. This result does not preclude the possibility that the tumor disappeared. However, owing to the low optical transparency of living tissue, the weak luminescence of tumor cells that had contracted to a small number may not have been detectable with IVIS.

As mentioned above, the correlation coefficient (r) between the scanner and caliper measurements is 0.96 in Fig. 2a and 0.727 in Fig. 4c. The only difference between Fig. 2a and 4c is the measurement day after CAR-T cell administration. The caliper measurements are performed on day 10 after CAR-T administration in Fig. 2a, whereas measurements are performed on endpoint (day 24) in Fig. 4. This implies that the tumor volume values measured using calipers or scanners appears to be correlated until approximately day 10 during the tumor growth phase. However, these differ significantly between the tumor regression phase and endpoint. To determine why this difference increases during the tumor regression phase, we performed a detailed comparison of each component of the tumor bulge measured using calipers or scanners at the endpoint (Fig. 5). Stereoscopic images of the tumor bulge on day 24 in the UTD and 1E6 CAR-T cell administration groups captured using a scanner are shown in Fig. 5a. The tumor bulge in the scanner is white, and the background tissue is blue. Fig. 5b shows a list of the components measured using a Vernier caliper or scanner for the same tumor bulge. It is noteworthy that although the differences in the longest diameter (major axis), shortest diameter (minor axis), and base area were not significantly different between the UTD and CAR-T cell groups, the height values were substantially different between the caliper and scanner measurements. (Fig. 5b). As mentioned above, the width of the tumor bulge was used rather than the height of the tumor bulge when calculating the tumor volume using calipers. As a result, the volume values differed substantially between the caliper and scanner. This indicates that the tumor bulge varied in height to a larger extent than in the area of its elliptical dome-shaped base, and regressed into a flat shape in response to variations in volume. This characteristic variation in tumor shape over time is considered to be the main reason for the large difference in tumor volume between the caliper and scanner measurements during the regression phase.

**Figure 5.**
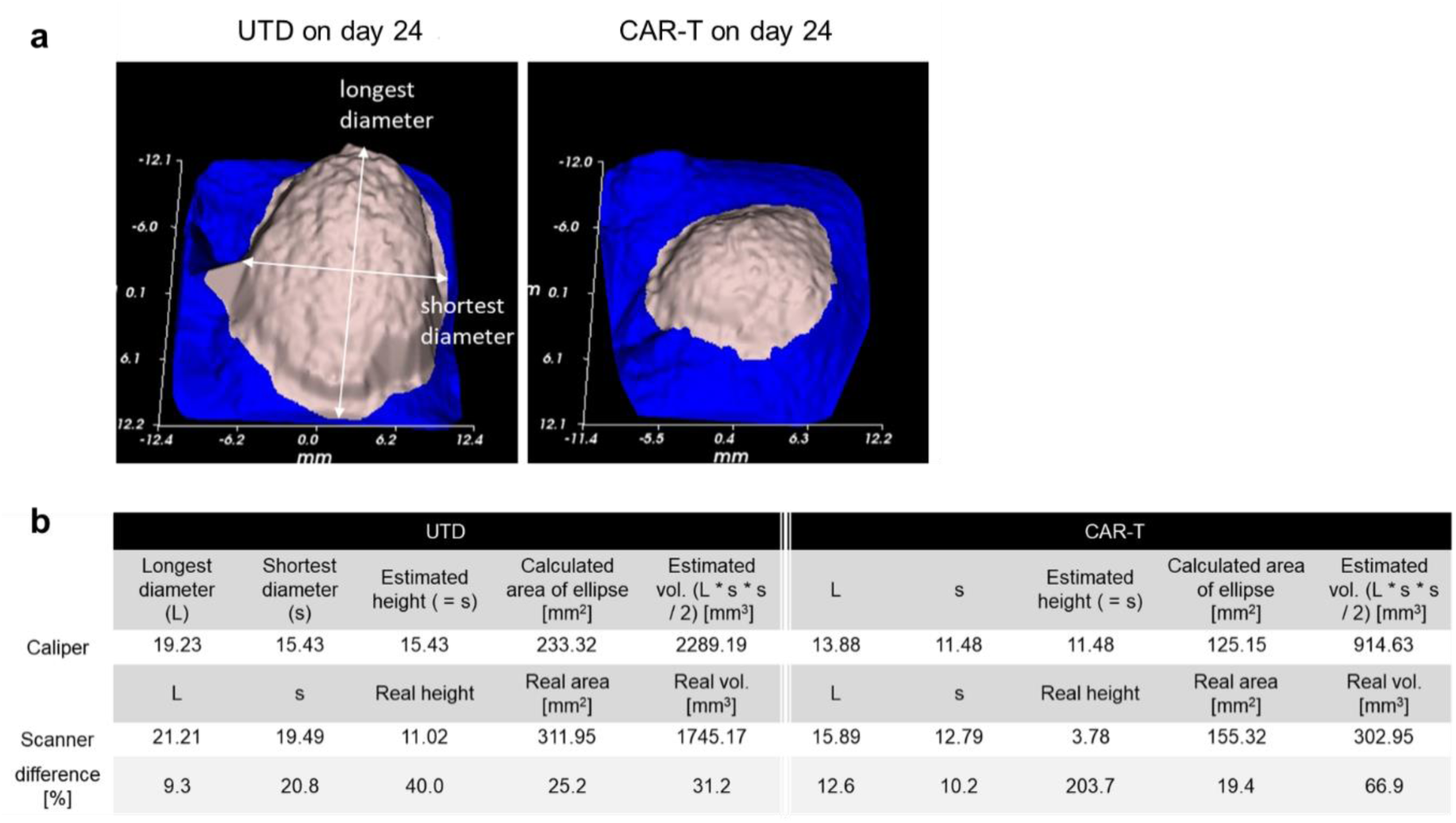
The characteristic shape variations during tumor regression resulted in differences in scanner and caliper measurements. (a) Example of 3D images captured using a 3D scanner. The left and right panels show the bulging of the tumor 24 days after the administration of UTD and CAR-T cells, respectively, to tumor-bearing mice. (b) The values measured using calipers or scanners for the above objects as shown in (a) are compared in detail, and the difference rates for each value are displayed.

## Discussion

In this paper, we report that 3D scanner measurements are accurate, have low variability, and are highly correlated with tumor weight. The scanner was observed to be more sensitive to volume regression than the caliper, particularly when the bulge flattened during tumor regression. Moreover, it showed drug efficacy with a higher resolution than a caliper. The accuracy of 3D scanners is effective for evaluating drug efficacy and ensuring product equivalence when changing raw materials or manufacturing processes during the development of cell therapy products.

Caliper measurement requires advance preparation such as practice for operation and repeated practice to match the subjective values to prevent inconsistencies in measurements by different operators. However, 3D scanner measurement is performed instantaneously and automatically. Therefore, no advance preparation is required. In addition, the scanner does not require training to administer luciferin intraperitoneally in advance for luminescence imaging, nor does it need to wait owing to the kinetics of luciferin. Therefore, 3D scanners can also contribute to the reduction of man-hours. In addition, scanner measurements are nondestructive and noninvasive, and it is unnecessary to administer chemicals such as luciferin for luminescence into the living body. Therefore, it can be measured intact. This is preferable because there is no concern regarding biotoxicity in nonclinical tests. Owing to the irregular shape of the subcutaneous tumor bulge and the heterogeneity of its internal tissue, improvements in optimal formulas and measurement methods have been proposed and discussed (Tomayko, et al., 1989; Jensen et al., 2008; Ayers et al., 2010; Hall et al., 2021). This study demonstrates that optical 3D scanning is simple and accurate with no interoperator variability. Moreover, it displays a higher correlation and linearity with tumor volume than caliper or IVIS measurements, particularly because of its high accuracy during the tumor regression phase.

Compared with the caliper measurement, the scanner measurement measured with exceptionally small differences between operators. Although the same test mouse groups were measured, significant differences existed between the operators when they measured with the calipers (Fig. 2). This result is considered to be largely owing to the subjective difference between operators regarding where the tumor bulge is located and where the width and length are located. This indicates the importance of practicing subject matching in advance. However, the scanner can measure the bulge volume only mechanically without subjective assessment. The scanner measurements revealed a higher discrimination capability over time than the caliper measurements. In particular, during the tumor regression phase, both the measurements showed the largest differences (Fig. 4). This could be owing to variations in tumor height (particularly during the tumor regression phase), which results in a flatter tumor shape and therefore, higher variability in caliper measurements (Figs.2, 4, and 5). This shape variation during tumor regression may have been caused by the tumor growing faster than destructively invading the host’s healthy subcutaneous tissue and by the overall tumor contracting faster than the subcutaneous tissue flattening owing to healing.

As shown in Fig. 4c, the correlation coefficient between the caliper and scanner measurements for day 24 was r = 0.727 and that between the weight and scanner measurements was almost equal at r = 0.726. However, because the correlation coefficient between the caliper and scanner for day 10 was r = 0.95 as shown in Fig. 2a, the caliper measurements for day 24 were more varied than those for day 10. These results indicated that the endpoint weight measurements were highly variable. Tissues from the tumor area were isolated in several autopsies. This may have caused variations owing to human differences. Therefore, it was inferred that the original true tumor weight and scanner measurements had correlation coefficients higher than r = 0.726. This in turn, indicates that the scanner measurements accurately represented values close to the true tumor volume.

Although the variability in tumor weight values should be noted, the optimal measurement range for the IVIS was estimated to be approximately 200–800 mg (Fig. 4b, c). When using IVIS in a drug efficacy study of subcutaneous tumor-bearing, it is preferable to adjust the amount of tumor bearing at the beginning of the study to vary within this weight range during the study. However, because of pseudo-progression (which causes CAR-T cells to accumulate in the tumor area) and necrosis caused by hypoxia, it is likely that there are fewer tumor cells than in the bulging volume (Chiou et al., 2015; Abrouk et al., 2018; Bertout et al., 2008). As shown in Fig. 4b, the tumor volume values for the 1E6 CAR-T group by caliper and scanner measurements peaked on day 11, whereas the IVIS values peaked earlier on day 5. Similarly, scanner values decreased to the initial values on day 24. However, the IVIS values decreased more rapidly to the initial values on day 19 and to almost zero on day 26. According to a previous study (Rice et al., 2000), the IVIS can detect luminescent signals in at least 500 subcutaneously transplanted cancer cells. Assuming that the average number of cells in a human weighing 70 kg is 37.2 trillion (Bianconi, et al., 2013), the average weight of one cell is approximately 1.88 ng, and 500 cells are estimated to weigh 940 ng. The minimum tumor weight was 106.4 mg when the IVIS value was zero as shown in Fig. 4c. This weight is larger than that of 500 cells. Therefore, it is assumed that the IVIS value decayed faster than the tumor weight value and fell below the detection sensitivity of our experimental system that used a conventional IVIS Lumina II machine. We need to consider the likelihood that the isolated tissues contained normal mouse epidermal cells. In the future, in order to accurately detect tumor cells with higher sensitivity, it is preferable to take images with longer exposure times, use more sensitive latest IVIS machine, and use the more sensitive luciferin / luciferases such as AkaBLI that would be able to detect even a single tumor cell implanted *in vivo* (Iwano, et al., 2018). High-sensitivity IVIS measurements would have been effective if the purpose of the study had been to track the complete disappearance of cancer cells at the single-cell level. This study focused on the superiority of scanners for comprehensive efficacy assessment accuracy from the perspective of CMC in the pharmaceutical industry. The most important thing is the capability to measure the transition with less variability and higher accuracy with high linearity throughout the efficacy study and to demonstrate that the values are statistically and significantly different from those of the control group, rather than to accurately determine the date when the tumor volume approaches zero. In this context, the scanner measurements were the most effective.

Note that when the IVIS measurement value is small, it may be difficult to assess whether the tumor volume has decreased or whether the intraperitoneal administration of the substrate has failed owing to operational error. In such a case, we need to re-administer the substrate and verify whether this low value is correct. However, this process depends on the subjectivity of each operator, which results in inaccurate values and increased man-hours for each measurement. To solve this problem, a fluorescent reagent such as IVISbrite D-luciferin ultra-bioluminescent substrate (Revvity Health Sciences Inc., USA) can be used to determine the success or failure of intraperitoneal administration by simultaneously measuring the luminescence and fluorescence imaging. We used it in this study. However, because the success of intraperitoneal injection is based on the relative fluorescence intensity, it was difficult to completely eliminate subjectivity. This operational problem appears to be one of the reasons why the IVIS measurement values varied throughout the drug efficacy test (Fig. 4c).

The advantages of 3D scanning over the calipers and IVIS measurements described in this paper are summarized in Table 2. All the types and models of commercially available 3D scanners are significantly more expensive than calipers and require initial investment. However, after being introduced, these provide many advantages. The accuracy is improved by eliminating the estimation and subjectivity in caliper measurements, errors caused by human manipulation in caliper and IVIS measurements, and differences between operators. In addition, no training is required to operate the scanner. The scanner can perform measurements rapidly without prior preparation or verification experiments and thereby, reduce man-hours. In addition, scanner measurements are safer because these do not require the administration of chemicals similar to those for IVIS measurements and can be performed intact. Again, as mentioned above, a robust measurement with a 3D scanner for efficacy evaluation results in improved accuracy, reduced man-hours, and improved safety. This contributes to the quality evaluation and characterization of products in the CMC of pharmaceutical companies.

**Table 2.** A summary of the advantages of optical 3D scanning compared with measurements using caliper or luminescence imaging.

|  | advantage of scanner<br>against caliper | advantage of scanner against<br>luminescence imaging |
| --- | --- | --- |
| **Improved**<br>**accuracy** | Elimination of estimated<br>values (Fig.2) | Reducing human interaction<br>(Discussion) |
|  | Elimination of subjectivity in<br>determining tumor shape<br>(Fig.2) | Elimination of human operations<br>such as laying the mouse in the<br>correct position (Table 1) |
|  | Reduction of human<br>operations such as mouse<br>holding (Table 1) | Elimination of inter-individual<br>differences in techniques such<br>as intraperitoneal administration<br>(Discussion) |
|  | Eliminating human differences<br>in technique (Fig.2) | The detection limit is large and<br>does not plateau (humane<br>endpoint is reached before the<br>detection limit) |
| **Reduced**<br>**man-hours** | No need for prior practice such<br>as subjective matching (Fig.2) | No need for advance<br>preparation such as Kinetics<br>test (Fig.4) |
| **Improved**<br>**safety** |  | Non-invasive and intact<br>measurement without<br>intraperitoneal administration |
|  |  | No chemical administration |

## Materials and Methods

### Animal studies

All the experiments and procedures involving animals conformed to the animal care and experimentation guidelines of the iPark Animal Experiments Committee and Institutional Animal Care and Use Committee (IACUC) (approval no. AU-00030666). NOD.Cg-*Prkdc*^scid^Il2rg^tm1Wjl^/SzJ (NSG) (The Jackson Laboratory, Inc.) female mice aged four–six weeks were used for the *in vivo* experiments. Five mice were housed in each cage.

### Murine subcutaneous xenograft models

GSU cells (RIKEN, JAPAN), a human cell line derived from gastric cancer, expressing Redfluc were generated using a lentiviral vector (PerkinElmer, Inc., USA). The cells were cultured in Dulbecco’s Eagle’s Medium (DMEM; Fujifilm Wako Chemicals, Corp., JAPAN) containing 10% fetal bovine serum (FBS; Biosera Inc., USA) and antibiotics, and subcultured by dissociation with trypsin (FUJIFILM Wako Pure Chemical Corp., JAPAN) every three days. Then, 5E6 (5 × 10^6) GSU-Redfluc cells were suspended in 100 uL DPBS (Dulbecco’s Phosphate Buffered Saline; Gibco Inc., USA) and xenografted subcutaneously into NSG mice. A week after tumor bearing, the luminescence intensity of the tumor was measured. The mice were grouped to average the amount of tumor carried by each mouse so that it was not biased from cage to cage. Thereafter, the different cell therapy products or PBS were administered intravenously to the mice via the tail vein in each cage. We used chimeric antigen receptor T (CAR-T) cells expressing armoring cytokines or untransduced (UTD) T cells prepared from human leukapheresis products (Cryopreserved Leukopak, Charles River Laboratories Cell Solutions, Inc., USA). The tumors were isolated and weighed at the study endpoint 35 days after inoculation.

### Tumor quantification

The tumor size was measured two times a week using calipers (Mitsutoyo Corp., JAPAN) or a 3D scanner (TM900, Peira, BELGIUM). Additionally, the luminescence and fluorescence intensity were measured one time a week using an IVIS (Lumina II, Caliper Life Sciences, Inc., USA) by intraperitoneal injection of 150 mg/kg luciferin, IVISbrite D-Luciferin Ultra Bioluminescent Substrate (Revvity Health Sciences Inc., USA), and analyzed using Living Image Software (version 4.7.3, Caliper Life Sciences, Inc., USA). Statistical analyses were performed using Microsoft Excel (Excel for Microsoft 365 MSO version 2310, Microsoft, Inc., USA) and R statistical software (R version 4.3.2. R Core Team (2023). R: A Language and environment for statistical computing. R Foundation for Statistical Computing, Vienna, Austria. https://www.R-project.org/). The Data in Fig. 2d were statistically analyzed using a one-way analysis of variance (ANOVA) followed by Bonferroni’s multiple comparisons. The data in Fig. 4b were statistically analyzed using an ANOVA followed by Dunnett’s multiple comparisons. The asterisks indicate statistically significant differences between the results of the UTD and various doses of CAR-T cells (*p < 0.1, **p < 0.05, and ***p < 0.01).

## Acknowledgement

Soichiro Ogaki and Satoko Hamanaka (Cell Therapy Technology and Product Engine, Research, Takeda Pharmaceutical Company) provided the CAR-T cells. This work was supported by Takeda Pharmaceutical Company.

## Author contributions

All authors were involved in drafting the article or revising it critically for important intellectual content, and all authors approved the final version to be published. T.K. designed the research and wrote the draft, perform the experiment and data analysis; M.K. performed the experiment; Y.N. performed the experiment and writing (review and editing), Y.T performed the experiment; H.A. performed the experiment and writing (review and editing), E.M. project administration and writing (review and editing).

## Competing Interest Statement

The authors are employees of Takeda Pharmaceutical Company.

## References

Teicher, B. A. Tumor models for efficacy determination. Mol. Cancer Ther. 5(10), 2435–43, DOI: 10.1158/1535-7163.MCT-06-0391 (2006 Oct).

Sausville, E. A. & Burger, A. M. Contributions of human tumor xenografts to anticancer drug development. Cancer Res. 66(7), 3351–4, DOI: 10.1158/0008-5472.CAN-05-3627 (2006 Apr 1).

Ruggeri, B. A., Camp, F. & Miknyoczki, S. Animal models of disease: pre-clinical animal models of cancer and their applications and utility in drug discovery. Biochem. Pharmacol. 87(1), 150– 61, DOI: 10.1016/j.bcp.2013.06.020. Epub 2013 Jun 28 (2014 Jan 1).

Liu, Y., Wu, W., Cai, C., Zhang, H., Shen, H. & Han, Y. Patient-derived xenograft models in cancer therapy: technologies and applications. Signal Transduct. Target Ther. 8(1), 160, DOI: 10.1038/s41392-023-01419-2 (2023 Apr 12).

Ovejera, A. A. Houchens, D. P. & Barker, A. D. Chemotherapy of human tumor xenografts in genetically athymic mice. Ann. Clin. Lab. Sci. 8, 50–56, (1978).

Hoffman, R. M. The multiple uses of fluorescent proteins to visualize cancer in vivo. Nat. Rev. Cancer. 5(10), 796–806, DOI: 10.1038/nrc1717 (2005 Oct).

de Jong, M., Essers, J. & van Weerden, W. M. Imaging preclinical tumour models: improving translational power. Nat. Rev. Cancer. 14(7), 481–93, DOI: 10.1038/nrc3751. Epub 2014 Jun 19 (2014 Jul).

Walsh, L. A. & Quail, D. F. Decoding the tumor microenvironment with spatial technologies. Nat. Immunol. 24(12), 1982–1993, DOI: 10.1038/s41590-023-01678-9. Epub 2023 Nov 27 (2023 Dec).

Sato, A., Klaunberg, B. & Tolwani, R. In vivo bioluminescence imaging. Comp. Med. 54(6), 631– 4, (2004 Dec).

Kelkar, M. & De, A. Bioluminescence based in vivo screening technologies. Curr. Opin. Pharmacol. 12(5), 592–600, DOI: 10.1016/j.coph.2012.07.014. Epub 2012 Sep 3 (2012 Oct).

Kato, T., et al. Human visual cortical function during photic stimulation monitoring by means of near-infrared spectroscopy. J. Cereb. Blood Flow Metab. 13(3), 516–520 (1993).

Smith, A. M., Mancini, M. C. & Nie, S. Bioimaging: second window for in vivo imaging. Nat. Nanotechnol. 4(11), 710–1, DOI: 10.1038/nnano.2009.326. PMID: 19898521; PMCID: PMC2862008 (2009 Nov).

Hong, G., Antaris, L. A. & Dai, H. Near-infrared fluorophores for biomedical imaging. Nat. Biomed. Eng. 1, 0010, 10.1038/s41551-016-0010 (2017).

Kobayashi, T., Haruta, M., Sasagawa, K., et al. Optical communication with brain cells by means of an implanted duplex micro-device with optogenetics and Ca2+ fluoroimaging. Sci. Rep. 6, 21247, 10.1038/srep21247 (2016).

Rice, B. W., Cable, M. D. & Nelson, M. B. In vivo imaging of light-emitting probes. J. Biomed. Opt. 6(4), 432–40, DOI: 10.1117/1.1413210 (2001 Oct).

Yeh, H. W., Karmach, O., Ji, A., Carter, D., Martins-Green, M. M. & Ai, H. W. Red-shifted luciferase-luciferin pairs for enhanced bioluminescence imaging. Nat. Methods. 14(10), 971–974, DOI: 10.1038/nmeth.4400. Epub 2017 Sep 4 (2017 Oct).

Endo, M. & Ozawa, T. Advanced Bioluminescence System for In Vivo Imaging with Brighter and Red-Shifted Light Emission. Int. J. Mol. Sci. 21(18), 6538, DOI: 10.3390/ijms21186538 (2020 Sep 7).

Cauchon, N. S., Oghamian, S., Hassanpour, S. & Abernathy, M. Innovation in Chemistry, Manufacturing, and Controls-A Regulatory Perspective From Industry. J. Pharm. Sci. 108(7), 2207–2237, DOI: 10.1016/j.xphs.2019.02.007. Epub 2019 Feb 19 (2019 Jul).

Girit, I. C., Jure-Kunkel, M. & McIntyre, K. W. A structured light-based system for scanning subcutaneous tumors in laboratory animals. Comp. Med. 58(3), 264–70, (2008 Jun).

Mu, P., Zhang, Z., Benelli, M., Karthaus, W. R., Hoover, E., Chen, C. C., Wongvipat, J., Ku, S. Y., Gao, D., Cao, Z., Shah, N., Adams, E. J., Abida, W., Watson, P. A., Prandi, D., Huang, C. H., de Stanchina, E., Lowe, S. W., Ellis, L., Beltran, H., Rubin, M. A., Goodrich, D. W., Demichelis, F. & Sawyers, C. L. SOX2 promotes lineage plasticity and antiandrogen resistance in TP53- and RB1-deficient prostate cancer. Science. 355(6320), 84–88, DOI: 10.1126/science.aah4307 (2017 Jan 6).

Delgado-SanMartin, J., Ehrhardt, B., Paczkowski, M., Hackett, S., Smith, A., Waraich, W., Klatzow, J., Zabair, A., Chabokdast, A., Rubio-Navarro, L., Rahi, A. & Wilson, Z. An innovative non-invasive technique for subcutaneous tumour measurements. PLoS One. 14(10), e0216690, DOI: 10.1371/journal.pone.0216690. PMID: 31609977; PMCID: PMC6791540 (2019 Oct 14).

Tomayko, M. M. & Reynolds, C. P. Determination of subcutaneous tumor size in athymic (nude) mice. Cancer Chemother. Pharmacol. 24, 148–154, 10.1007/BF00300234 (1989).

Jensen, M. M., Jørgensen, J. T., Binderup, T. & Kjaer, A. Tumor volume in subcutaneous mouse xenografts measured by microCT is more accurate and reproducible than determined by 18F-FDG-microPET or external caliper. BMC Med. Imaging. 8, 16, DOI: 10.1186/1471-2342-8-16. PMID: 18925932; PMCID: PMC2575188 (2008 Oct 16).

Ayers, G. D., McKinley, E. T., Zhao, P., Fritz, J. M., Metry, R. E., Deal, B. C., Adlerz, K. M., Coffey, R. J. & Manning, H. C. Volume of preclinical xenograft tumors is more accurately assessed by ultrasound imaging than manual caliper measurements. J. Ultrasound Med. 29(6), 891–901, DOI: 10.7863/jum.2010.29.6.891. PMID: 20498463; PMCID: PMC2925269 (2010 Jun).

Hall, C., von Grabowiecki, Y., Pearce, S. P., Dive, C., Bagley, S. & Muller, P. A. J. iRFP (near-infrared fluorescent protein) imaging of subcutaneous and deep tissue tumours in mice highlights differences between imaging platforms. Cancer Cell Int. 21(1), 247, DOI: 10.1186/s12935-021-01918-8. PMID: 33941186; PMCID: PMC8091726 (2021 May 3).

Carmeliet, P. & Jain, R. K. Principles and mechanisms of vessel normalization for cancer and other angiogenic diseases. Nat. Rev. Drug Discov. 10(6), 417–27, DOI: 10.1038/nrd3455. PMID: 21629292 (2011 Jun).

Bertout, A. J., Patel, A. S. & Simon, C. M. The impact of O2 availability on human cancer. Nat. Rev. Cancer. 8(12), 967–75, DOI: 10.1038/nrc2540. Epub 2008 Nov 6 (2008 Dec).

Chiou, V. L. & Burotto, M. Pseudoprogression and immune-related response in solid tumors. J. Clin. Oncol. 33, 3541–3543, (2015).

Abrouk, N., Oronsky, B., Caroen, S., Ning, S., Knox, S. & Peehl, D. A note on improved statistical approaches to account for pseudoprogression. Cancer Chemother. Pharmacol. 81, 621– 626, (2018).

Bianconi, E., Piovesan, A., Facchin, F., Beraudi, A., Casadei, R., Frabetti, F., Vitale, L., Pelleri Chiara, M., Tassani, S., Piva, F., Perez-Amodio, S., Strippoli, P. & Canaider, S. An estimation of the number of cells in the human body. Ann. Hum. Biol. 40(6), 463–71, DOI: 10.3109/03014460.2013.807878. Epub 2013 Jul 5 (2013 Nov–Dec).

Iwano, S., Sugiyama, M., Hama, H., Watakabe, A., Hasegawa, N., Kuchimaru, T., Tanaka, K. Z., Takahashi, M., Ishida, Y., Hata, J., Shimozono, S., Namiki, K., Fukano, T., Kiyama, M., Okano, H., Kizaka-Kondoh, S., McHugh, T. J., Yamamori, T., Hioki, H., Maki, S. & Miyawaki, A. Single-cell bioluminescence imaging of deep tissue in freely moving animals. Science. 359(6378), 935–939, DOI: 10.1126/science.aaq1067. PMID: 29472486 (2018 Feb 23).

